# Machine Learning Gap-Fills Missing Transporter Kinetics in Biosystems Across Scales

**DOI:** 10.64898/2026.07.02.735998

**Authors:** Sizhe Qiu, Zhangyu Guo, Weiming Tu, Yingping Zhuang, Shengbo Wu, Guan Wang

**Author notes:** Corresponding authors (S. Wu), (G. Wang).

## Abstract

Understanding transporter kinetics is essential for deciphering metabolite exchanges in biosystems, particularly for cells subject to substrate gradients. Nevertheless, the prediction of transporter kinetic parameters, maximum rate per gram protein (V_max_) and Michaelis-Menten constant (K_m_), has not yet been tackled. Here, we developed the first compound-protein interaction machine learning model of transporter V_max_ and K_m_, MMTKPred, which achieved R^2^=0.553, RMSE=1.155 mmol/hr/g Protein and R^2^=0.330, RMSE=0.935 mM for log10-scaled V_max_ and K_m_ prediction, respectively. Moreover, we demonstrated MMTKPred’s predictive power across biosystem scales, from capturing transporter kinetics modulated by point mutations and substrate changes at the molecular level, to enabling substrate-sensitive metabolic modelling of non-model yeasts at the cellular level, and rationalizing inter-species substrate competition in co-cultures. Collectively, MMTKPred effectively models metabolite transport spanning from molecular to multi- species scales, thereby offering a computational tool for rational microbial cell factory optimization.

**Graphical abstract:** 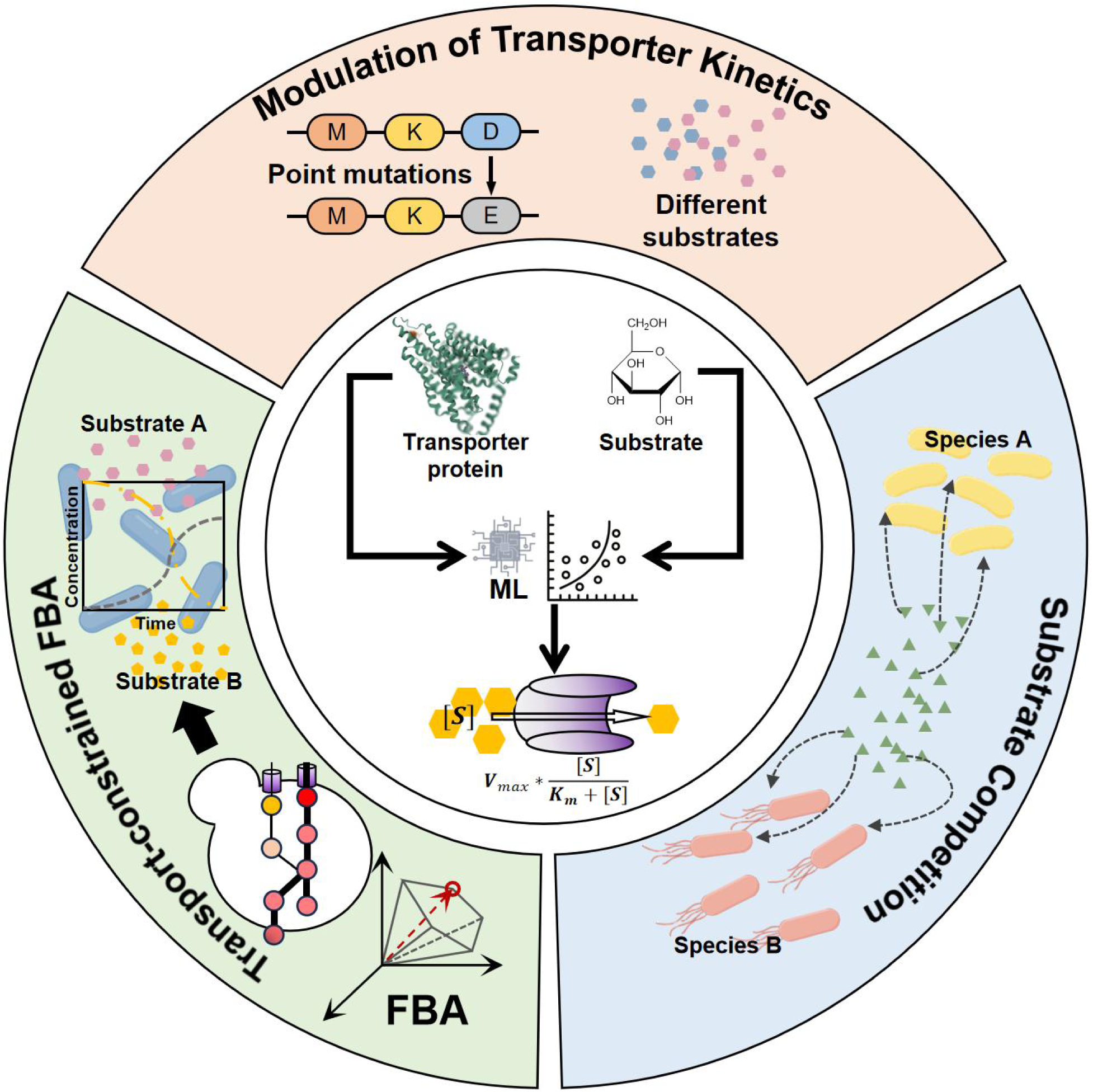

**Highlights:**

1. MMTKPred, first transporter kinetics CPI model, reaches ∼1 log10 RMSE for V_max_ and K_m_.
2. MMTKPred captures the effects of point mutations and substrate changes on transporters.
3. Predicted kinetics enables substrate sensitivity in metabolic flux modelling.
4. Predicted kinetics explains inter-species substrate competition outcomes.

## 1. Introduction

Transporter proteins (e.g., channels and carriers) largely dominate metabolite flux across cellular membranes, connecting cells to their environments, and hence, transporter kinetics is essential for decoding metabolite exchanges in biosystems across scales ^1,2^. In industrial bioprocesses, transporter kinetics provides a quantitative basis for determining substrate uptake and product export rates in chassis cells, which are key operation parameters for bioreactors with substrate gradients ^3^. For microbial cell factories, inefficient transporters often become the bottlenecks of productivity, and hence, researchers adopted various transporter engineering strategies, including directed evolution and rational structural design, to boost bio-manufacturing efficiency by enhancing substrate uptake, intracellular metabolite transfer, and product export ^4,5^. Beyond individual microbial cell factories, transporter kinetic traits, shaped by evolutionary adaptation, directly impact inter-species metabolic interactions and consequently structure microbial consortia ^6,7^. Therefore, transporter kinetics mechanisitically bridges physiological functions with the dynamics of natural and engineered biosystems.

In recent years, machine learning (ML) models for transporter-substrate specificity have progressed from class-level annotation to specific substrate prediction. For example, MONSTROUS employs graph neural networks to annotate 12 major transporters ^8^, and SPOT uses a transformer architecture to predict transporter–substrate pairs ^9^. However, no predictor has yet been constructed to estimate transporter kinetic parameters, V_max_ (maximum rate per unit protein) and K_m_ (Michaelis–Menten constant), unlike the development of compound-protein interaction (CPI) models of enzyme kinetic parameters ^10^. Given the aforementioned importance of transporter kinetics and its data scarcity in public biochemical databases (e.g., UniProt ^11^) ^12^, this shortfall critically hinders the quantitative analysis of metabolic fluxes in single/multi- species systems and the rational engineering of transporters for enhanced bioproduction ^5^.

With the aim to accurately predict transporter kinetics, this study endeavored to develop the first CPI model of transporter V_max_ and K_m_, namely MMTKPred, using pre-trained language models of proteins and compounds, and ML regression models like XGBoost. After model performance evaluation, MMTKPred was used to predict the effects of point mutations and substrate changes on transporter kinetics, constrain substrate-specific transportation capacity in metabolic modelling, and mechanistically explain outcomes of inter-species substrate competition. Collectively, this study established MMTKPred as a powerful computational tool for modelling and engineering metabolite transport in biosystems.

## 2. Methods

### 2.1 Datasets

To construct CPI models for transporter V_max_ and K_m_, this study collected all data entries containing the UniProt ID of transporter protein, substrate name, organism information, EC number, and the target value (V_max_ or K_m_) from UniProt ^11^, BRENDA ^13^, and SABIO-RK ^14^ in February, 2026. The datasets included only transporters that do not modify substrates during transport and that exhibit Michaelis-Menten kinetics. This study assumed that most transporter kinetics can be reasonably approximated using the Michaelis-Menten equation ^15^. Phosphotransferase systems (PTSs) were excluded as they phosphorylate substrates, and do not belong to EC 7 but EC 2.7.1.-, which has already been well-characterized by predictors of enzyme kinetic parameters ^10^. Transporter protein sequences were queried using UniProt IDs, and SMILES strings of substrates were queried from PubChem ^16^. The units of all V_max_ values were unified as 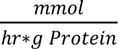, and the units of all K_m_ values were unified as *mM*. After the entries from UniProt, BRENDA, and SABIO-RK were merged, all redundant entries were removed. For entries with the same protein sequences and SMILES strings but different V_max_ values, only the entry with the largest V_max_ was kept. For entries with the same protein sequences and SMILES strings but different K_m_ values, the geometric mean of these K_m_ values was calculated. For SMILES strings with disconnected components, only the larger component was kept. Entries with SMILES strings containing metal ions were all removed. In the end, the datasets of V_max_ and K_m_ had 691 and 1706 unique entries, respectively.

For both V_max_ and K_m_, to ensure that the test datasets remained unseen, the training and test datasets were split randomly while controlling for sequence similarity, such that no similar sequences appeared in both sets. All protein sequences were clustered with a similarity cut-off of 50% using MMseqs2 ^17^, and then, these clusters were randomly split in a 2:8 ratio. The training and test datasets of V_max_ contained 559 and 132 entries, respectively, and the training and test datasets of K_m_ contained 1431 and 275 entries, respectively (**Figures S1&S2 in SI**).

### 2.2 Sequence and molecular feature computation

The sequence embeddings of transporter proteins were computed by ESM-2 (esm2_t33_650M_UR50D) ^18^, and the molecular embeddings of substrates were computed by ChemBERTa ^19^ from SMILES strings (**Figure 1**). Mean and max pooling were performed on sequence and molecular embeddings to generate Seq Mean/Max and Mol Mean/Max, which were considered as non-engineered features (Features (No FE), FE: feature engineering) in this study. For feature engineering, principal component analysis (PCA) with 256 components was used to generate dimensionality-reduced (DR) features: Seq Mean (DR), Seq Max (DR), Mol Mean (DR), and Mol Max (DR). Subsequently, element-wise multiplication of Seq Mean (DR) and Mol Mean (DR) produced the CxP Mean feature, and element-wise multiplication of Seq Max (DR) and Mol Max (DR) produced the CxP Max feature. Seq Mean/Max (DR), Mol Mean/Max (DR), and CxP Mean/Max were considered as engineered features (Features (FE)). Besides, Morgan fingerprints (MFPs) with 1024 bits were computed from substrate SMILES strings using RDKit ^20^ as an additional feature.

**Figure 1.**
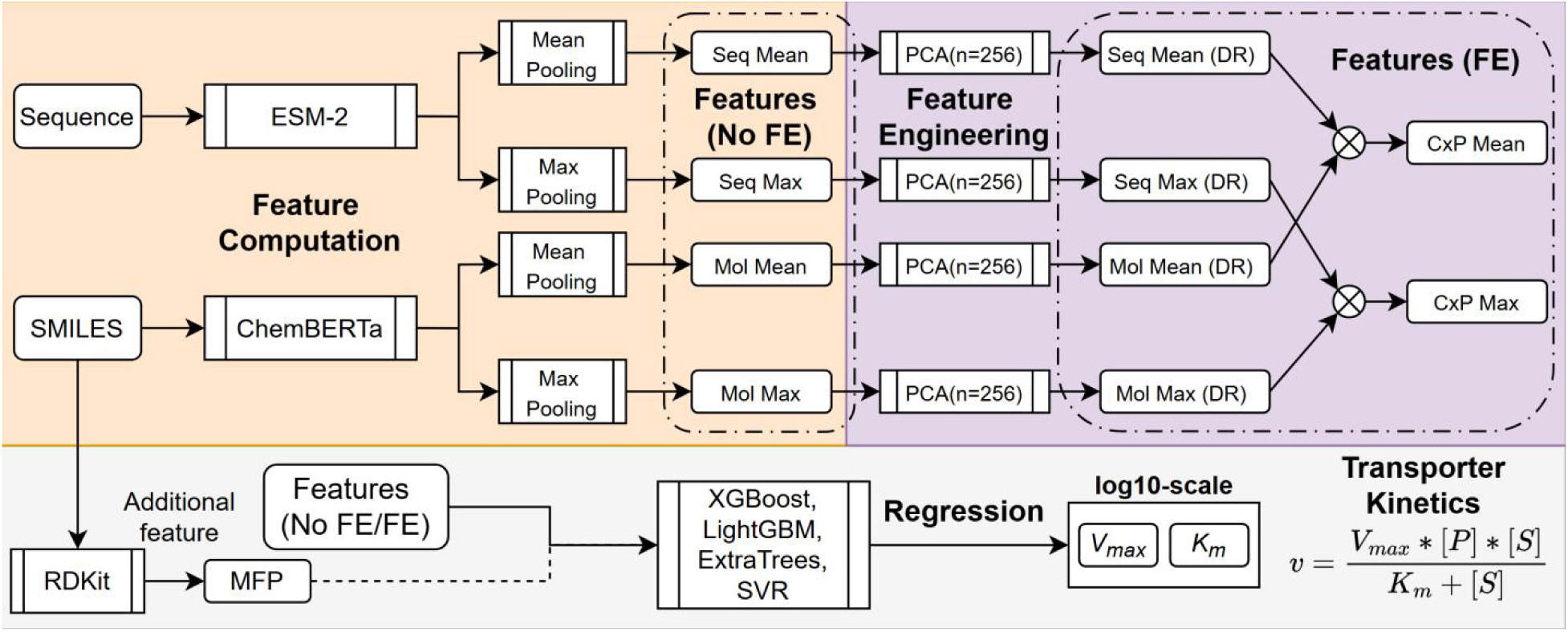
Model diagram of MMTKPred for transporter V_max_ and K_m_ prediction. Seq Mean: mean-pooled sequence embedding, Seq Max: max-pooled sequence embedding, Mol Mean: mean-pooled molecular embedding, Mol Max: max-pooled molecular embedding, DR: dimension reduced, CxP Mean: element-wise product of Seq Mean and Mol Mean, CxP Max: element-wise product of Seq Max and Mol Max, FE: feature engineering, NoFE: without feature engineering, MFP: Morgan fingerprint, V_max_: maximum rate per gram protein, K_m_: Michaelis-Menten constant, [P]: transporter protein mass per gram DCW, [S]: substrate concentration.

### 2.3 Machine learning model construction and hyperparameter optimization

To predict transporter V_max_ and K_m_, this study trialled 4 regression models that were commonly used in CPI models or sequence-based models of protein properties: XGBoost, LightGBM, ExtraTrees, and SVR ^21,22,23^ (**Figure 1**). Also, 4 different combinations of features were examined for model training and evaluation: (1) Features (No FE), (2) Features (No FE) + MFP, (3) Features (FE), and (4) Features (FE) + MFP (**Section 2.2**). To note, V_max_ and K_m_ were predicted separately, and the target values were all log10-scaled. Hyperparameter optimization of different machine learning models was conducted by Optuna ^24^, and the search spaces of hyperparameters were detailed in **Table S1 in SI**. With V_max_ and K_m_ predicted, the Michaelis-Menten transporter kinetics can be estimated as:

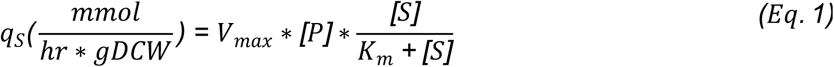

*q_S_* is the cell-specific transporter rate for a certain substrate, *[P]* is the transporter protein mass per gram dry cellular weight (DCW), and *[S]* is the substrate concentration. For trained fine-tuned machine learning models of transporter V_max_ and K_m_, feature importance analysis was performed by computing the SHAP value ^25^ for each feature.

### 2.4 Flux balance analysis constrained by predicted transporter kinetics

Conventional flux balance analysis (FBA) computes metabolic fluxes by optimizing an objective function (the default is growth rate, *μ*) (Eq.2) with steady-state mass conservation (Eq.3) ^26^. *A* Is the stoichiometric matrix, and *ν* is the vector of reaction fluxes. The integration of predicted transporter kinetic parameters (Eq.1) into FBA constrains the transportation capacity (exchange fluxes) and enables the sensitivity to substrate identities and concentrations, giving rise to the transport-constrained FBA (TCFBA) framework. As the total mass of membrane proteins is limited, the summation of transporter protein mass has an upper bound, *C* (Eq.4 & Eq.5). 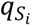 is the exchange flux of the substrate *S_i_*, [*P_i_*] is the mass per DCW of the transporter protein *P_i_*.

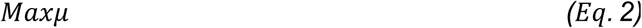

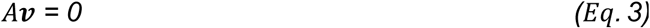

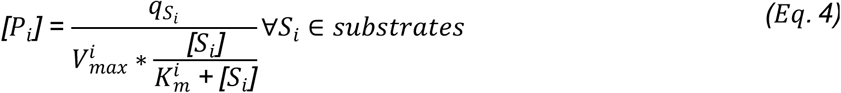

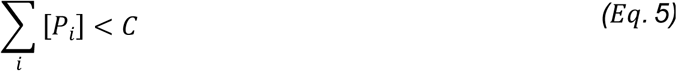

The sensitivity of TCFBA to substrate concentrations also facilitates dynamic FBA ^27^, as it can approximate the change of exchange fluxes based on substrate concentrations.

### 2.5 Metagenomic data processing for inter-species substrate competition

In this study, the experimental inter-species competition outcomes for different substrates were obtained from Kehe et al., 2021 ^7^, and the metagenomic sequences were requested from the same source. 101 unique inter-species substrate competitions, spanning 11 bacterial species and 9 substrates were selected, excluding all substrates that can be transported via PTSs, such as fructose. In the dataset of Kehe et al., 2021, the effect score of bacteria 1 on bacteria 2 is quantified as log2-scale difference of growth yields of bacteria 2 in co-culture with bacteria 1 and mono-culture, and thus, the more negative “1 on 2” effect score indicates a stronger substrate competition ability. For transporter protein identification, Pyrodigal 3.7.1 ^28^ was used to annotate protein sequences from metagenomic sequences, and then substrate-specific transporters were identified by sequence alignment against TCDB database ^29^ using Diamond ^30^. For Diamond version 2.1.25, the max hit number was set as 5, and e-value threshold was set as 1e-5, other parameters were set as default values. The average similarity between annotated transporter sequences from 11 bacterial species and the sequences in the training datasets was below 40% (**Figure S9 in SI**). The substrate information was mapped to each matched hit using Transporter Classification IDs in TCDB database. All annotated transporter sequences of these 11 bacterial species were deposited in bugcu_filtered_proteins.tsv at https://github.com/SizheQiu/MMTKPred/data/competition/.

## 3. Results

### 3.1 XGBoost achieves high accuracy in transporter V_max_ prediction

Among 4 machine learning models and different feature sets, XGBoost had superior performance in log10 V_max_ prediction, and feature engineering did not improve the prediction accuracy (**Figure 2A**). The top 2 models were XGBoost using Features (No FE) (R^2^=0.553, RMSE=1.155 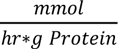, MAE=0.788 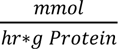) and XGBoost using Features (No FE)+MFP (R^2^=0.546, RMSE=1.164 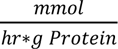, MAE=0.814 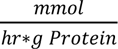) (**Figure 2BC**). The optimized hyperparameter sets of these 2 models were detailed in **Table S2 in SI**. As there has not been any transporter V_max_ predictor published yet, this study did not perform a model performance comparison here. SHAP-based feature importance analysis of XGBoost using Features (No FE) and XGBoost using Features (No FE)+MFP both showed that the top important features were mainly mean- and max-pooled sequence features, suggesting that the regression of V_max_ relied primarily on protein sequence features (**Figure 2DE**). By contrast, MFP did not appear in top important features of XGBoost using Features (No FE)+MFP (**Figure 2E**), and the distributions of dimension-reduced Features (No FE)+MFP also showed that MFP contributed little to discriminate between low and high V_max_ values (**Figure S3 in SI**). The performance evaluation for different value ranges demonstrated that XGBoost using Features (No FE) performed slightly better at high value range (top 25%), while XGBoost using Features (No FE)+MFP performed slightly better at low value range (lower 25%) (**Figure 2F**). Furthermore, this study found that XGBoost using Features (No FE) could capture the evolutionary adaptation whereby microorganisms (*S. cerevisiae* and *E. coli*) have significantly higher transporter Vmax than mammalian cells (*H. sapiens* and *M. musculus*) (**Figure 2GH**), as microorganisms are subject to much stronger nutrient competition ^31^. In short, XGBoost using Features (No FE) was recognized as the best model of transporter V_max_ and it captured the evolutionary insight of cross-species transporter V_max_ variation.

**Figure 2.**
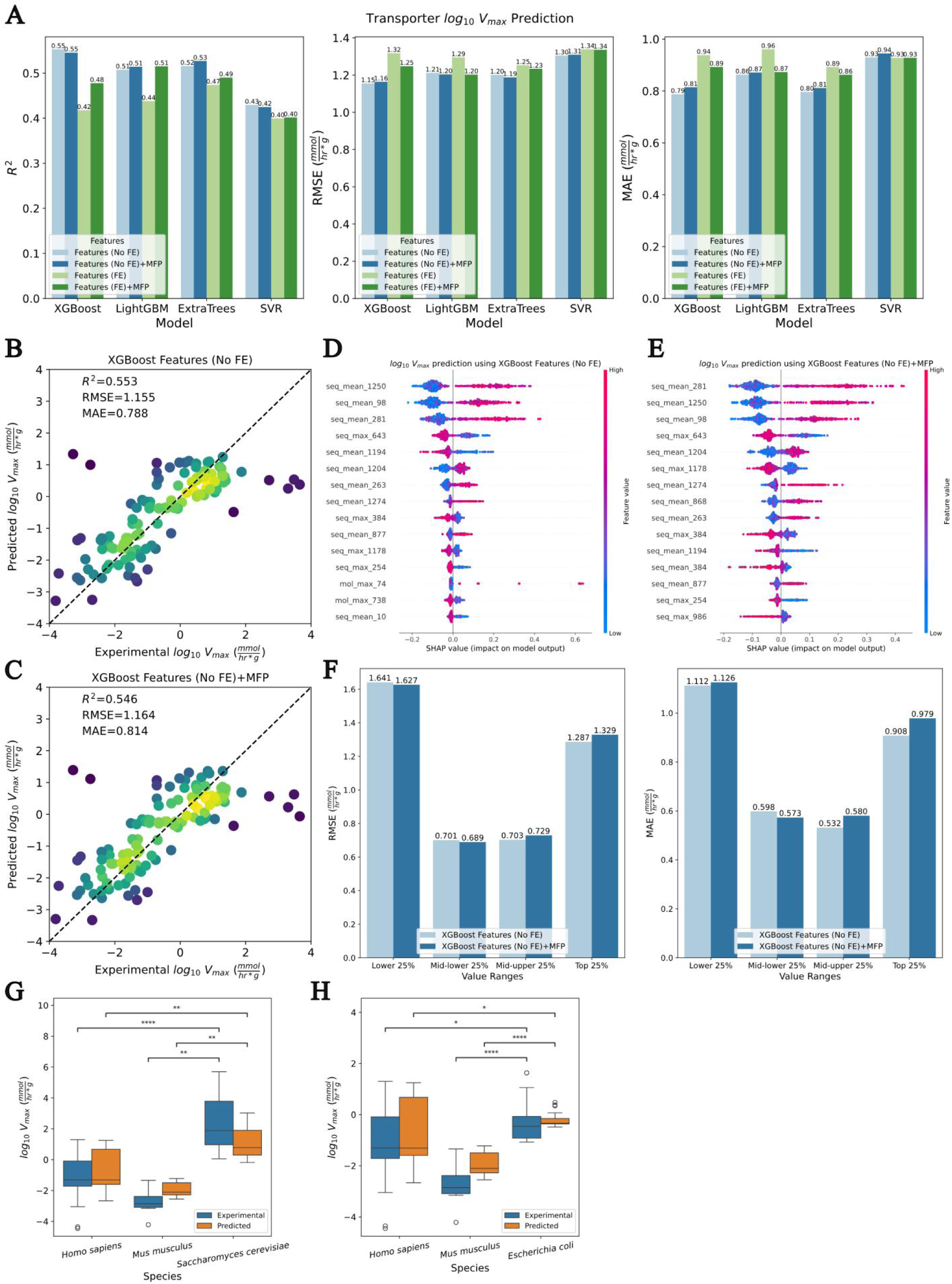
Model performance evaluation for transporter V_max_ prediction. (A) R^2^, RMSE, MAE of XGBoost, LightGBM, ExtraTrees, and SVR using Features (No FE), Features (No FE)+MFP, Features (FE), and Features (FE)+MFP. FE: feature engineering, NoFE: without feature engineering, MFP: morgan fingerprint. (B) Experimental log10 V_max_ and predicted log10 V_max_ by XGBoost using Features (No FE) (R^2^=0.553, RMSE=1.155 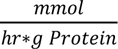, MAE=0.788 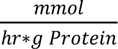). (C) Experimental log10 V_max_ and predicted log10 V_max_ by XGBoost using Features (No FE)+MFP (R^2^=0.546, RMSE=1.164 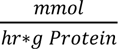, MAE=0.814 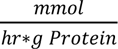). (D) SHAP-based feature importance analysis of XGBoost using Features (No FE). (E) SHAP-based feature importance analysis of XGBoost using Features (No FE)+MFP. (F) RMSEs and MAEs of XGBoost using Features (No FE) and XGBoost using Features (No FE)+MFPs for transporter log10 V_max_ in different value ranges (lower 25%, middle-lower 25%, middle-upper 25%, top 25%). (G) Experimental and XGBoost-predicted (Features (No FE)) transporter log10 V_max_ values for *Homo sapiens*, *Mus musculus*, and *Saccharomyces cerevisiae*. (H) Experimental and XGBoost-predicted (Features (No FE)) transporter log10 V_max_ values for *Homo sapiens*, *Mus musculus*, and *Escherichia coli*.

### 3.2 Feature engineering allows gradient boosting models to outperform existing models in transporter K_m_ prediction

In contrast to transporter log10 V_max_ prediction, feature engineering boosted the prediction performances in transporter log10 K_m_ prediction, and XGBoost using Features (FE) and LightGBM using Features (FE) were found as top 2 models (**Figure 3A**). Meanwhile, the weak discriminative ability of dimension-reduced Features (No FE) for K_m_ reaffirmed the necessity of feature engineering (**Figure S4 in SI**). XGBoost using Features (FE) had an accuracy of R^2^=0.330, RMSE=0.935 *mM*, MAE=0.727 *m M*, and LightGBM using Features (FE) had an accuracy of R^2^=0.325, RMSE=0.939 *mM*, MAE=0.727 *mM* (**Figure 3BC**). The optimized hyperparameter sets of these 2 models were detailed in **Table S2 in SI**. Regarding feature importance analysis, the presence of “CxP_mean” in top important features of XGBoost using Features (FE) showed that element-wise multiplication of sequence and molecular features (**Section 2.2**) obtained useful information for transporter-substrate interactions (**Figure 3D**), while top important features of LightGBM using Features (FE) only contained dimension-reduced sequence and molecular embeddings (**Figure 3E**). In the performance comparison of previously published K_m_ predictors and top 2 models of transporter K_m_ in this study, the top 2 models outperformed UniKP ^32^, CatPred ^33^, EITLEM ^34^, and KinForm ^23^ (**Figure 3F**). Subsequently, this study evaluated the performances of the top 2 models and CatPred for different value ranges of log10 K_m_, and found that XGBoost using Features (FE) and LightGBM using Features (FE) had lower prediction errors than CatPred at middle 50% and lower 25% ranges (**Figure 3G**). With respect to cross-species transporter K_m_, the experimental K_m_ values in the test dataset did not show significant cross-species differences (**Figure S5 in SI**). To sum up, XGBoost using Features (FE) reached a superior accuracy in transporter K_m_ prediction.

**Figure 3.**
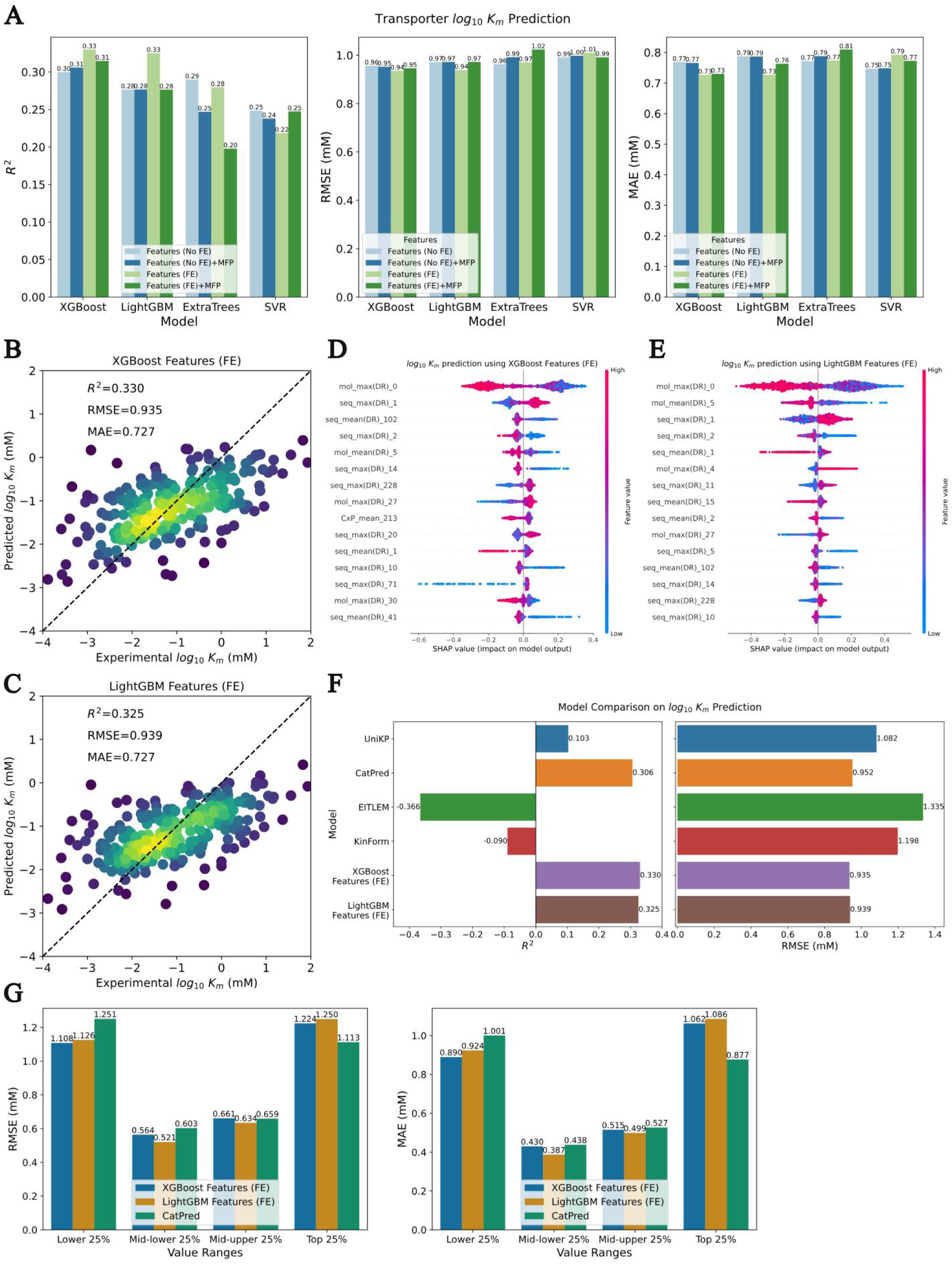
Model performance evaluation for transporter K_m_ prediction. (A) R^2^, RMSE, MAE of XGBoost, LightGBM, ExtraTrees, and SVR using Features (No FE), Features (No FE)+MFP, Features (FE), and Features (FE)+MFP. FE: feature engineering, NoFE: without feature engineering, MFP: morgan fingerprint. (B) Experimental log10 K_m_ and predicted log10 K_m_ by XGBoost using Features (FE) (R^2^=0.330, RMSE=0.935 *mM*, MAE=0.727 *mM*). (C) Experimental log10 K_m_ and predicted log10 K_m_ by LightGBM using Features (FE) (R^2^=0.325, RMSE=0.939 *mM*, MAE=0.727 *mM*). (D) SHAP-based feature importance analysis of XGBoost using Features (FE). (E) SHAP-based feature importance analysis of LightGBM using Features (FE). (F) R^2^ and RMSE of UniKP, CatPred, EITLEM, KinForm, XGBoost using Features (FE), and LightGBM using Features (FE). (G) RMSEs and MAEs of XGBoost using Features (FE), LightGBM using Features (FE), and CatPred for transporter log10 K_m_ in different value ranges.

### 3.3 Predicted transporter kinetics captures the effects of point mutations and substrate changes

After the performance evaluation of MMTKPred revealing prediction errors close to one order of magnitude for both V_max_ and K_m_, this study next assessed its extrapolation capability for mutated transporter proteins and varied substrates. We first employed MMTKPred to predict the effects of point mutations on transporter efficiency for 3 different transporters: (1) facilitated glucose transporter member 9 (SLC2A9) from *H. sapiens* ^35^, (2) multidrug resistance ABC transporter (BmrA) from *Bacillus subtilis* ^36^, (3) phenylalanine permease (PheP) of *E. coli* ^37^ (**Figure 4A-D**). All V_max_ predictions were conducted by XGBoost using Features (No FE) (**Sections 3.1**). As the maximum transportation efficiency of SLC2A9, BmrA, and PheP were not quantified in the same or convertible unit as V_max_ in this study, we could only qualitatively assess whether the model predicted the trends of V_max_ changes. For the three single-point mutations on SLC2A9, the model accurately predicted the decrease of V_max_ by I335V, W110A, and W110F (**Figure 4A**). The reduction of BmrA transportation activity by E474D, E474Q, and E474F mutations was consistent in experimental data and prediction results (**Figure 4B**). The predicted V_max_ of 33 PheP mutants had significant positive correlation with the experimental relative activity (**Figure 4C**). Regarding proline-to-alanine substitutions on 16 conserved proline residues of PheP, the trends of predicted V_max_ and experimental relative activity were highly consistent, particularly for the substantial reduction in activity caused by the P341A substitution (**Figure 4D**). In general, the model could account for the effects of single-point mutations on transporter efficiency.

**Figure 4.**
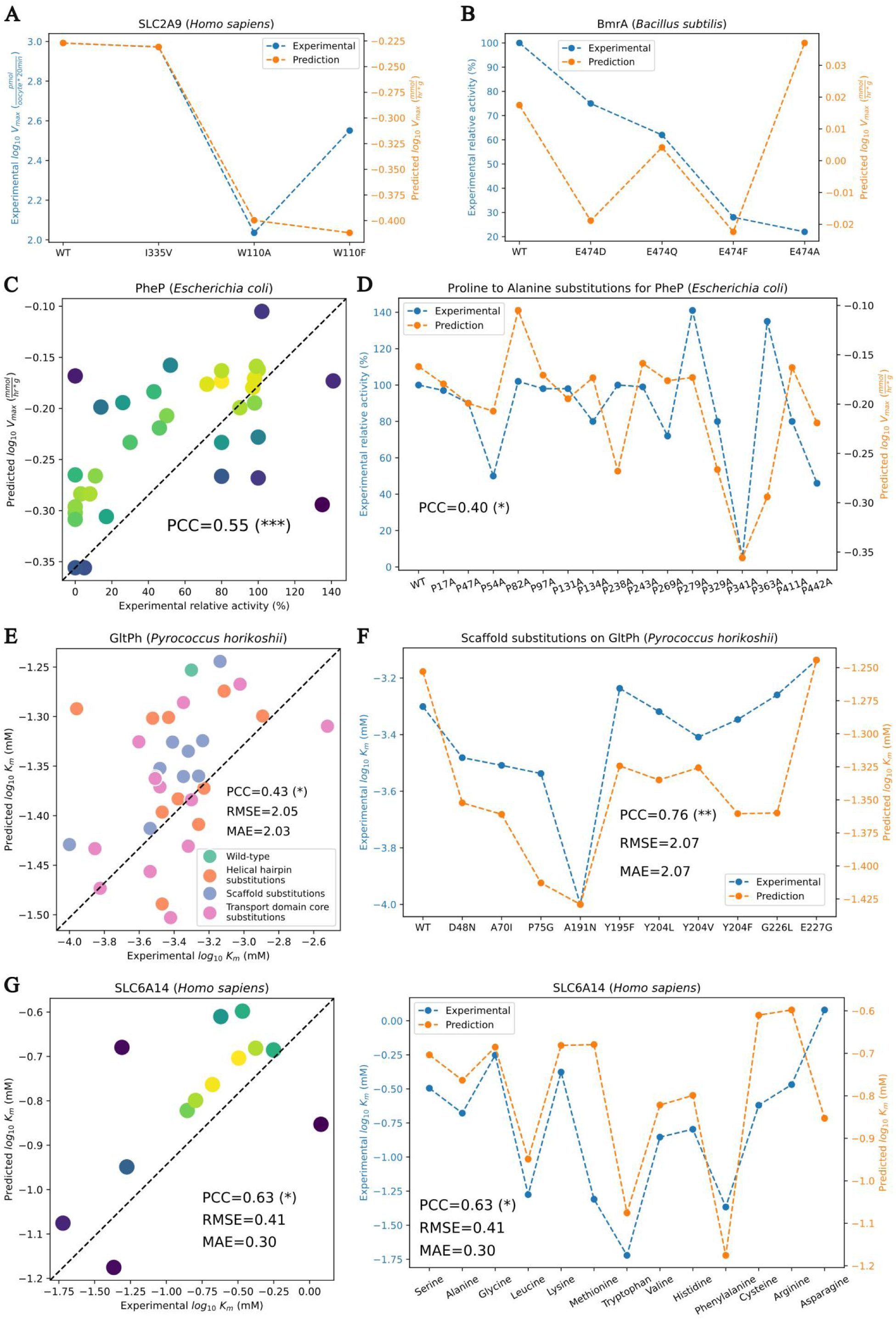
Predicting the effects of point mutations and substrates on transporter V_max_ and K_m_. (A) Experimental and predicted log10 V_max_ of wild-type and mutated SLC2A9. The unit of experimental V_max_ is pmol/(oocyte*20min). SLC2A9: facilitated glucose transporter member 9 from *H. sapiens*. (B) Experimental relative activities (%) and predicted log10 V_max_ of wild-type and mutated BmrA. BmrA: multidrug resistance ABC transporter from *B. subtilis*. (C) Experimental relative activities (%) and predicted log10 V_max_ of wild-type and mutated PheP. PheP: phenylalanine permease of *E. coli*, PCC: Pearson correlation coefficient. (D) Experimental relative activities (%) and predicted log10 V_max_ of PheP with proline to alanine substitutions. (E) Experimental and predicted log10 K_m_ of wild-type and mutated GltPh. GltPh: sodium-aspartate symporter from *Pyrococcus horikoshii*. (F) Experimental and predicted log10 K_m_ of GltPh with single-point mutations on the scaffold region. (G) Experimental and predicted log10 K_m_ of SLC6A14 for different amino acids. SLC6A14: sodium/chloride-dependent amino acid transporter from *H. sapiens*. Blue line: experimental value, Orange line: predicted value, *: p-value<0.05, **: p-value<0.01, ***: p-value<0.001.

Regarding transporter K_m_, we used MMTKPred to predict the effects of point mutations on sodium-aspartate symporter (GltPh) from *Pyrococcus horikoshii* ^38^ and substrate changes on sodium/chloride-dependent amino acid transporter (SLC6A14) from *H. sapiens* ^39^, respectively (**Figure 4E-G**). All K_m_ predictions were conducted by XGBoost using Features (FE) (**Sections 3.2**). The predicted K_m_ of GltPh with point mutations on helical hairpin, scaffold, and transport domain core had significant positive correlation with experimental K_m_, although the numerical error remained large (log10 RMSE>2) (**Figure 4E**). Compared to other two regions, the model had higher accuracy for K_m_ changes caused by mutations on the scaffold region (**Figure 4E, Figures S6&S7 in SI**). For 13 different amino acids transported by SLC6A14, the model accurately predicted substrate-specific K_m_ (log10 RMSE=0.41), and identified leucine, tryptophan, and phenylalanine as substrates with top affinities (**Figure 4G**). On the whole, MMTKPred could effectively predict the modulation of transporter kinetics by point mutations and substrate changes, empowering high-throughput *in silico* screening for candidate transporter proteins and nutrient sources to optimize microbial cell factories.

### 3.4 Predicted transporter kinetics enables substrate-sensitive metabolic modelling

Following the validation of MMTKPred’s predictive power on transporter kinetics modulation by single point mutations and substrate changes, this study then evaluated its applicability to metabolic flux modelling. While *Yarrowia lipolytica* and *Kluyveromyces marxianus*, two widely-used non-model yeasts, lack quantified kinetic data for both enzymes and transporters ^40^, transport-constrained FBA (TCFBA) coupled with MMTKPred-predicted transporter kinetics (**Section 2.4**) tackles this limitation, allowing low- cost simulation of their growth and metabolism on different carbon sources (**Figure 5**). The genome-scale metabolic models used were iYali v4.1.2 ^41^ for *Y. lipolytica* and iSM996 ^42^ for *K. marxianus*. Considering the prediction errors of transporter V_max_ and K_m_ (**Sections 3.1&3.2**), a loose upper bound of membrane transporter proteins was applied, 0.5 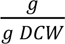^43^. First, transporter V_max_ and K_m_ of different carbon sources were predicted using information in **Table S3 in SI**, as required by subsequent case studies (**Figure S8 in SI**). The growth rates of *Y. lipolytica* and *K. marxianus* using different carbon sources were predicted and compared to experimental biomass yields, obtained from Mukhtar et al., 2018 ^44^ and Goshima et al., 2013 ^45^ (**Figure 5AB**). The predicted growth rates of *Y. lipolytica* by static TCFBA quantitatively reflected that glucose and maltose were preferred carbon sources for growth quantified by biomass yields (**Figure 5A**). Static TCFBA also accurately identified glucose, galactose, and mannose as carbon sources optimal for *K. marxianus* growth, but underestimated the growth rates of *K. marxianus* using sucrose (**Figure 5B**). In short, TCFBA coupled with MMTKPred- predicted transporter kinetics accurately determines substrate- specific growth kinetics.

**Figure 5.**
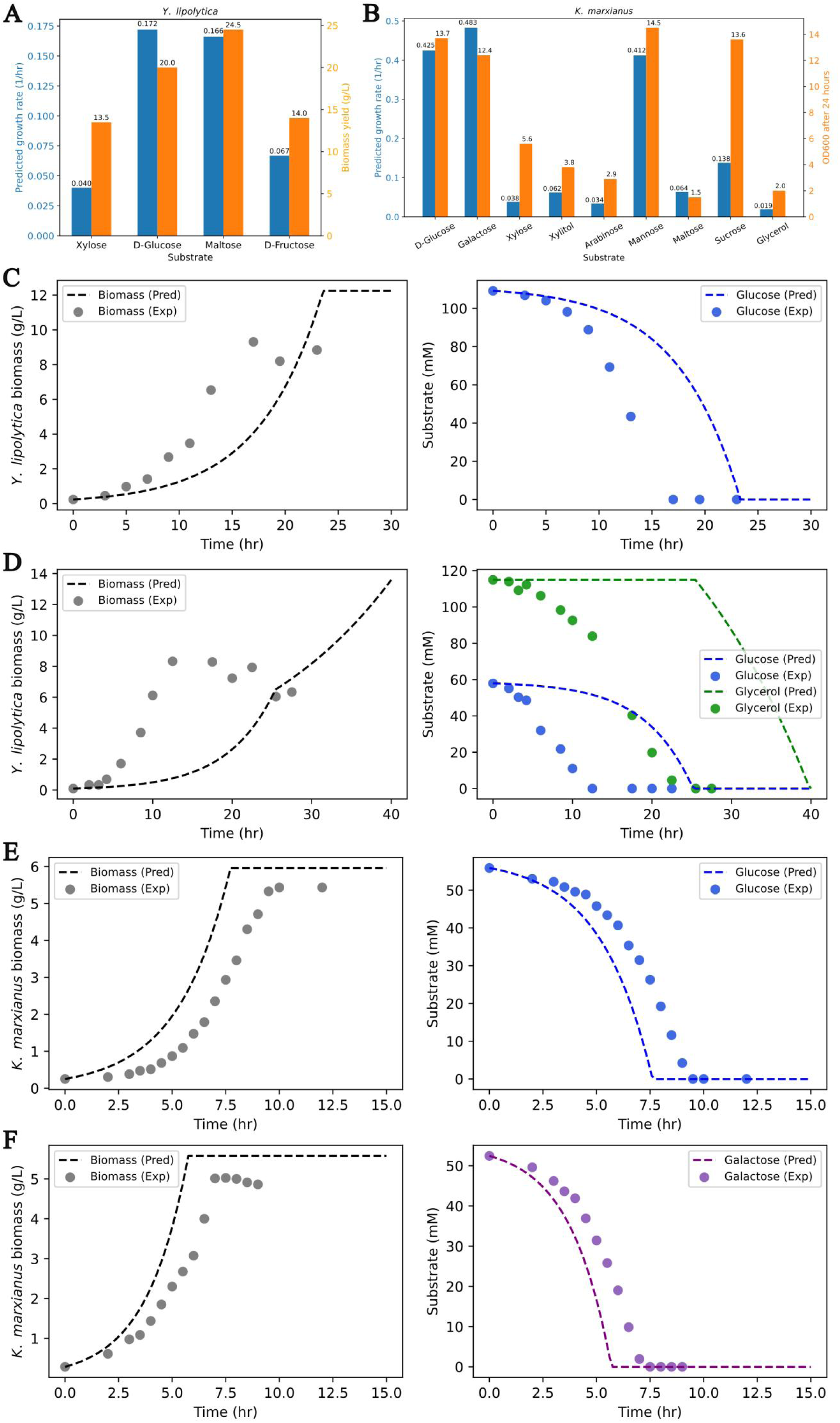
Transport-constrained metabolic modelling of *Y. lipolytica* and *K. marxianus* utilizing different carbon sources. (A) Predicted growth rates and experimental biomass yields of *Y. lipolytica* using xylose, glucose, maltose and fructose. (B) Predicted growth rates and OD600 after 24 hours of *K. marxianus* using glucose, galactose, xylose, xylitol, arabinose, mannose, sucrose, and glycerol. (C) Predicted and experimental biomass accumulation and substrate consumption for *Y. lipolytica* using glucose. (D) Predicted and experimental biomass accumulation and substrate consumption for *Y. lipolytica* using both glucose and glycerol. (E) Predicted and experimental biomass accumulation and substrate consumption for *K. marxianus* using glucose. (F) Predicted and experimental biomass accumulation and substrate consumption for *K. marxianus* using galactose. Exp: experimental value, Pred: predicted value.

Next, dynamic TCFBA was performed for (1) *Y. lipolytica* utilizing glucose and glycerol ^46^ and (2) *K. marxianus* utilizing glucose, galactose, and lactose ^47^. The transition from exponential to stationary phases of *Y. lipolytica*, due to glucose depletion, were quantitatively characterized by dynamic TCFBA, but the underestimated glucose consumption rate resulted in the deviation between predicted and experimental growth curves (**Figure 5C**). Dynamic TCFBA simulated diauxic growth during the co- metabolism of glucose and glycerol, although the utilization rate of glycerol was substantially underestimated (**Figure 5D, Figure S9A in SI**). For *K. marxianus*, the growth and substrate consumption curves for galactose and glucose utilization were simulated and exhibited high consistency with experimental measurements (**Figure 5EF**), but the utilization rate of lactose was underestimated (**Figure S9B in SI**). The prediction errors for *Y. lipolytica* on glycerol and *K. marxianus* on lactose (**Figure S9 in SI**) were most likely caused by the under-representation of these two substrates in the training datasets of transporter V_max_ and K_m_. In brief, TCFBA with predicted substrate-specific constraints on exchange fluxes adequately enables sensitivity to heterogeneous nutrient conditions in constraint-based metabolic modelling.

### 3.5 Predicted transporter kinetics explains inter-species substrate competition outcomes

Beyond individual microbial cell factories, this study also explored the predictive power of MMTKPred in co- culture systems. Given that MMTKPred showed the capability to capture cross-species V_max_ variation due to evolutionary adaptation (**Section 3.1**), this study attempted to use predicted transporter kinetic parameters (V_max_ and K_m_) to explain inter-species substrate competition outcomes (**Section 2.5**). The sum of predicted V_max_/K_m_ for all annotated substrate-specific transporter proteins in metagenomic sequences of a bacterial species was considered as that species’ predicted substrate competition score, and that score showed a significant negative correlation with the measured inter-species effect score (**Figure 6A**). Predicted transporter V_max_/K_m_ values explained 53.5% of inter-species substrate competition outcomes, and could achieve high accuracy for competitions involving most bacterial species (e.g., *Citrobacter freundii*, *Buttiauxella izardii*) and substrates (e.g., alanine, serine) (**Figure 6BC**). For example, the model accurately predicted the outcomes of 18 out of 27 inter-species competitions for alanine (**Figure 6C, Figure S10 in SI**). If excluding competitions for glycerol, which is absent in the training datasets of MMTKPred, the accuracy of competition outcome prediction could reach 61.4%. The analysis of experimental and predicted substrate competition victory rates for all substrates, and specifically for alanine and glutamine, with highest numbers of competitions (**Figure S10 in SI**), showed that the competitiveness of *Pseudomonas koreensis* was largely overestimated, while that of *Klebsiella aerogenes* and *Enterobacter ludwigii* were underestimated, which were major error sources (**Figure 6D**). Overall, MMTKPred- predicted transporter kinetics mechanistically explains inter- species substrate competition outcomes, aiding mixed- species culture control and thereby contributing to rational multi- species cell factory design.

**Figure 6.**
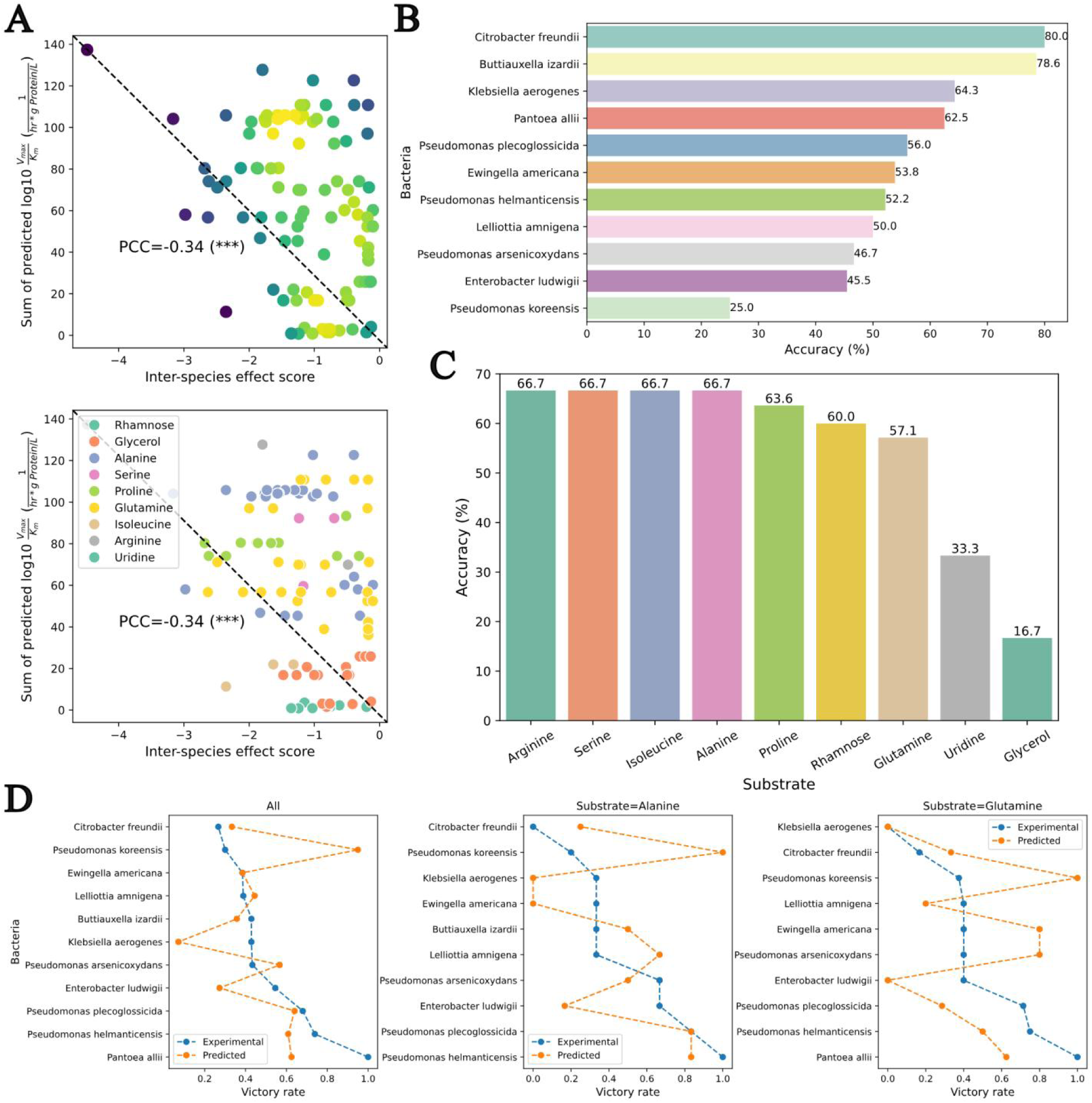
Prediction of inter-species substrate competition outcomes selected from Kehe et al. ^7^ spanning 11 bacterial species and 9 carbon sources with estimated transporter V_max_ and K_m_. (A) The correlation between sums of predicted log10 V_max_/K_m_ and inter-species effect scores for different substrates (p-value<1e-3). (B) The prediction accuracy of each species’ nutrient competition outcomes. (C) The prediction accuracy of inter-species substrate competition outcomes of each substrate. (D) Experimental and predicted substrate competition victory rates of each species for all substrates, just alanine, and just glutamine. ***: p-value<0.001.

## 4. Discussion

Transporter kinetics is essential for quantitative analysis, modelling, and engineering of metabolite transport in biosystems across different scales, from single transporter machinery to inter-species metabolic interactions. Given the importance of transporter kinetics and the scarcity of experimental data, a predictive model of transporter V_max_ and K_m_ is required to address this knowledge gap, which has hitherto remained unexplored. Hence, this study employed ESM-2 and ChemBERTa to encode transporter sequences and substrate molecules, respectively (**Section 2.2**), and trialled 4 different regression models to build the first predictor of transporter V_max_ and K_m_, MMTKPred (**Section 2.3**). Model evaluation identified XGBoost as the best regression model for both V_max_ (R^2^=0.553, RMSE=1.155 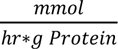, MAE=0.788 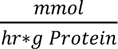 and K_m_ (R^2^=0.330, RMSE=0.935 *mM*, MAE=0.727 *mM*), and engineering of sequence and molecular features promoted the superior performance of MMTKPred over other K_m_ predictors (**Sections 3.1&3.2**). Furthermore, MMTKPred could capture the evolutionary insight into cross- species variation in transporter V_max_ (**Section 3.1**).

Following model evaluation, the study harnessed MMTKPred to predict the modulation of transporter kinetics via point mutations and substrate changes, showcasing its potential as a computational tool for transporter selection and engineering toward microbial cell factory optimization ^4^ (**Section 3.3**). With respect to metabolic modelling, TCFBA with predicted transporter kinetics gap-filled the protein cost of exchange fluxes and enabled the sensitivity to substrate identities and levels (**Section 3.4**), which has been missing in most enzyme-constrained metabolic models ^48^, and thus contributed to the development of metabolic digital twins for microbial cell factories under heterogeneous nutrient conditions. Moreover, employing predicted transporter kinetics to mechanistically model inter- species substrate competition extended quantitative metabolite transport modelling to co- culture level (**Section 3.5**), suggesting that MMTKPred could be used to design control strategies for multi-species cell factories ^49,50^.

Despite the achievements discussed above, there remain several limitations in MMTKPred that hinder the prediction accuracy of transporter kinetics. The prediction errors of V_max_ and K_m_ are both close to log10 RMSE of 1, and hence the errors of Eq.1 and V_max_/K_m_ are theoretically log10 RMSE of 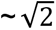 due to error propagation, yielding inaccuracy in dynamic metabolic modelling (**Section 3.4**) and inter-species substrate competition prediction (**Section 3.5**). The major bottleneck limiting prediction accuracy is the small size of the training dataset, compared with 16249 entries in the DLTKcat dataset ^51^, and the imbalanced distributions of V_max_ and K_m_ values (**Figures S1&S2 in SI**). Oversampling and loss reweighting algorithms (e.g., SMOTE ^52^) can mitigate the imbalance, but the number of entries in rare value regions of the MMTKPred training datasets is too small for these algorithms to take effect (**Figures S1&S2 in SI**). Additionally, inaccuracies in experimental data curated from databases can also impair model performance ^53^. Therefore, using high-throughput transporter assays to generate measured V_max_ and K_m_ for rare value regions, along with strict data quality control, is necessary to further improve the prediction accuracy of transporter kinetics. Another shortcoming is the neglect of environmental factors (e.g., pH) that affect transporter protein conformation and, consequently, transport kinetics ^54^. Including transporter assay metadata could help account for environmental factors, however, most transporter assay results curated from databases currently lack such information. The assumption of idealized Michaelis- Menten kinetics also limits MMTKPred’s applicability, as some transporters exhibit complex mechanisms (e.g., Hill kinetics) ^55^, but the scarcity of relevant training data currently precludes their incorporation. As expected, addressing the limitations outlined here would significantly improve the performance of MMTKPred in future research.

Taken together, this study produced the first CPI model of transporter kinetic parameters, V_max_ and K_m_, with prediction errors close to one order of magnitude, and demonstrated its predictive power in biosystems across scales, shedding light on metabolic digital twins of chassis cells under heterogeneous nutrient conditions and precise exchange flux control for rational strain engineering.

## Acknowledgements

This research was funded by the National Key Research and Development Program of China (Grant No. 2024YFA0920300), National Key Research and Development Program of China, (Grant No. 2024YFA0920200), Natural Science Foundation of China (Grant No. 22578122), the Taishan Scholars Program of Shandong Province (Grant No. tspn202408281), Natural Science Foundation of Shanghai (Grant No. 25ZR1402110), the Explorers Program of Shanghai (Basic Research Funding) (Grant No. 25TS1401100), Shanghai Rising-Star Program (Grant No. 21QA1402400), and the Jiangsu Provincial Major Science and Technology Special Project (Grant No. BG2025044). Additionally, I would like to thank Milan Sobota of University of Oxford for his assistance with the metagenomic data processing.

## Author contributions

Dr. Sizhe Qiu: Conceptualization, Data curation, Formal analysis, Investigation, Methodology, Visualization, Writing-original draft; Zhangyu Guo: Data curation, Visualization; Prof. Weiming Tu: Writing-review & editing; Prof. Yingping Zhuang: Project administration, Supervision, Writing-review & editing; Prof. Shengbo Wu: Project administration, Supervision, Writing-review & editing; Prof. Guan Wang: Project administration, Supervision, Writing-review & editing.

## Data availability statement

The code and data are openly available at https://github.com/SizheQiu/MMTKPred and supplementary information.

## Abbreviations

CPI: compound-protein interaction
DCW: dry cellular weight
FBA: flux balance analysis
FE: feature engineering
K_m_: Michaelis-Menten constant
MAE: mean absolute error
ML: machine learning
PCC: Pearson correlation coefficient
PTS: phosphotransferase system
R^2^: r-squared, the coefficient of determination
RMSE: root mean squared error
V_max_: maximum rate per gram protein

