## Supplementary material for "Machine Learning Gap-Fills Missing Transporter Kinetics in Biosystems Across Scales": Figures S1-S10, Tables S1-S3

### Supplementary Information

#### 1. Supplementary Figures

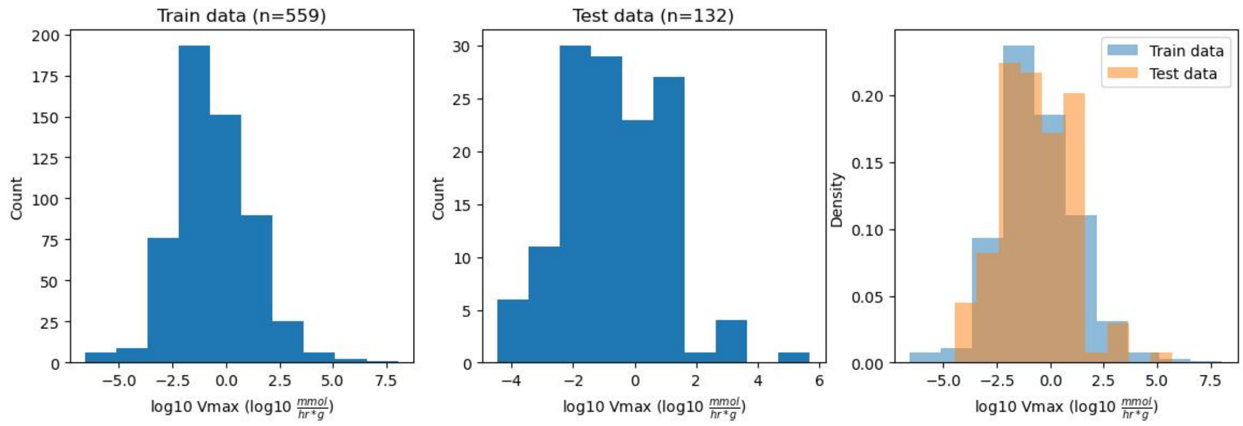


Figure S1. Distribution of log10 $V_{max}$ values in train and test datasets.

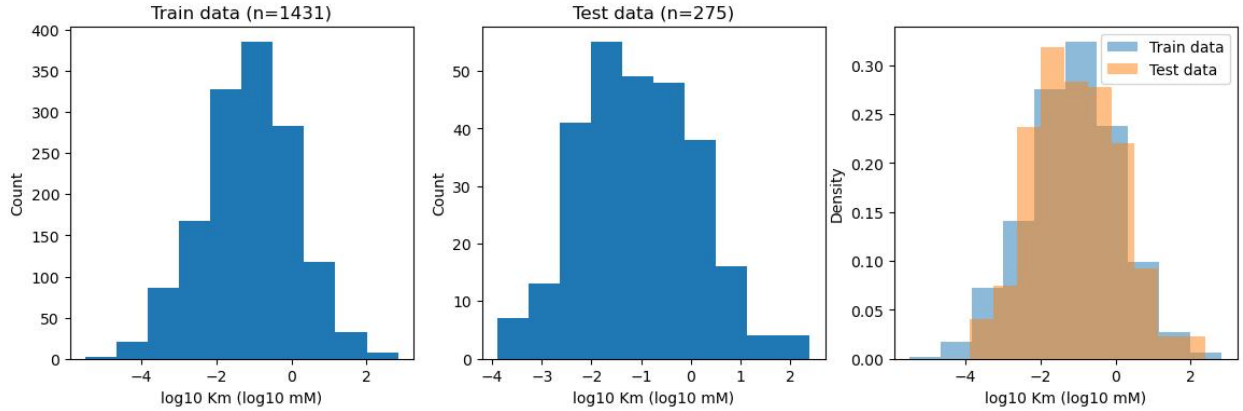


Figure S2. Distribution of log10 $K_m$ values in train and test datasets.

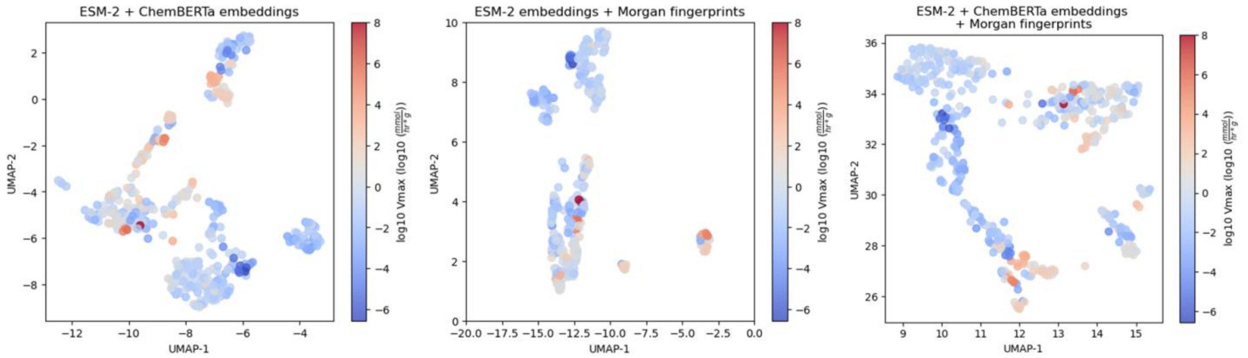


Figure S3. UMAP of (1) max- and mean-pooled sequence and molecular embeddings, (2) max- and mean-pooled sequence embeddings and morgan fingerprints, (3) max- and mean-pooled sequence and molecular embeddings, and morgan fingerprints for 559 entries in the train dataset of $V_{max}$.

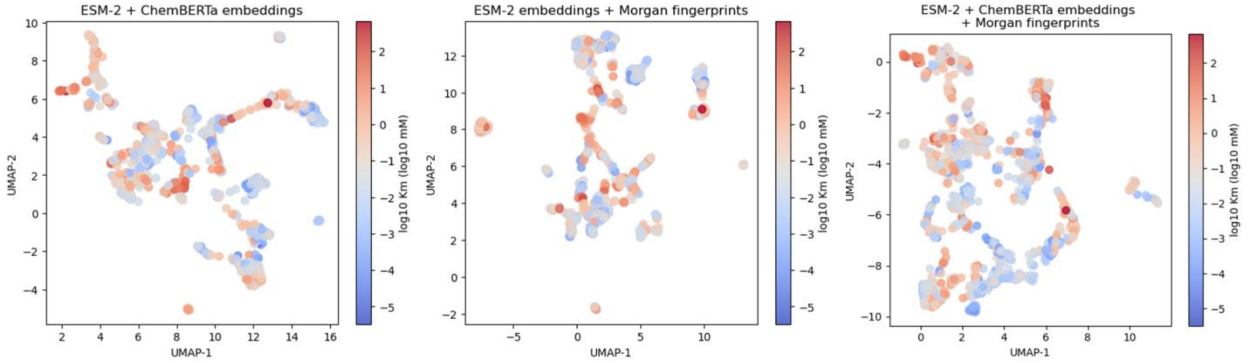


Figure S4. UMAP of (1) max- and mean-pooled sequence and molecular embeddings, (2) max- and mean-pooled sequence embeddings and morgan fingerprints, (3) max- and mean-pooled sequence and molecular embeddings, and morgan fingerprints for 1431 entries in the train dataset of $K_m$.

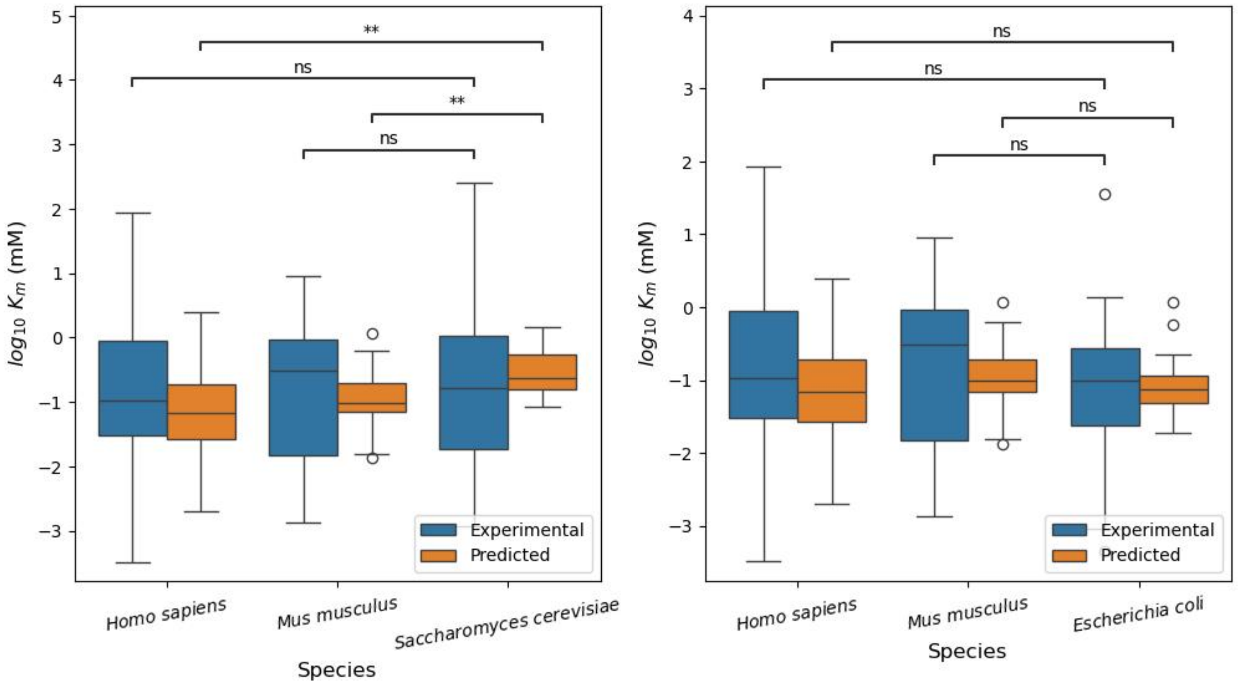


Figure S5. (Left) Experimental and XGBoost-predicted (Features (FE)) transporter log10 $K_m$ values for *Homo sapiens*, *Mus musculus*, and *Saccharomyces cerevisiae*. (Right) Experimental and XGBoost-predicted (Features (FE)) transporter log10 $K_m$ values for *Homo sapiens*, *Mus musculus*, and *Escherichia coli*.

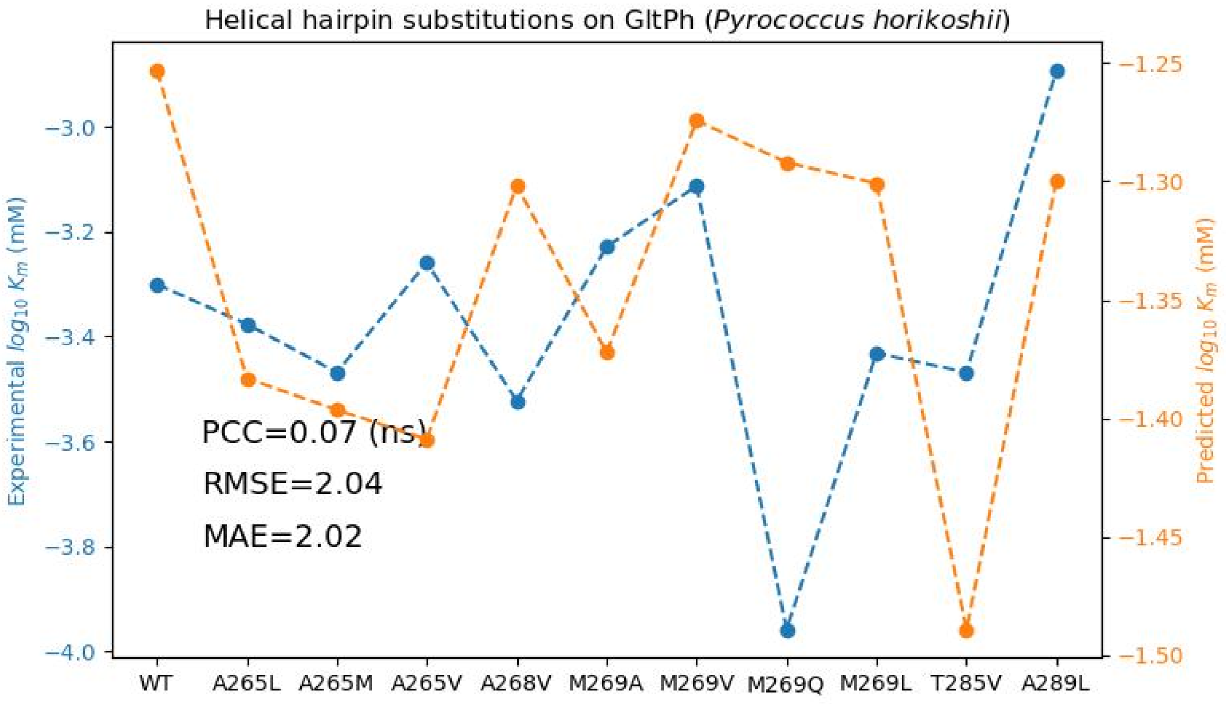

Figure S6. Experimental and predicted log10 $K_m$ of wild-type GltPh and GltPh with single-point mutations on the helical hairpin.

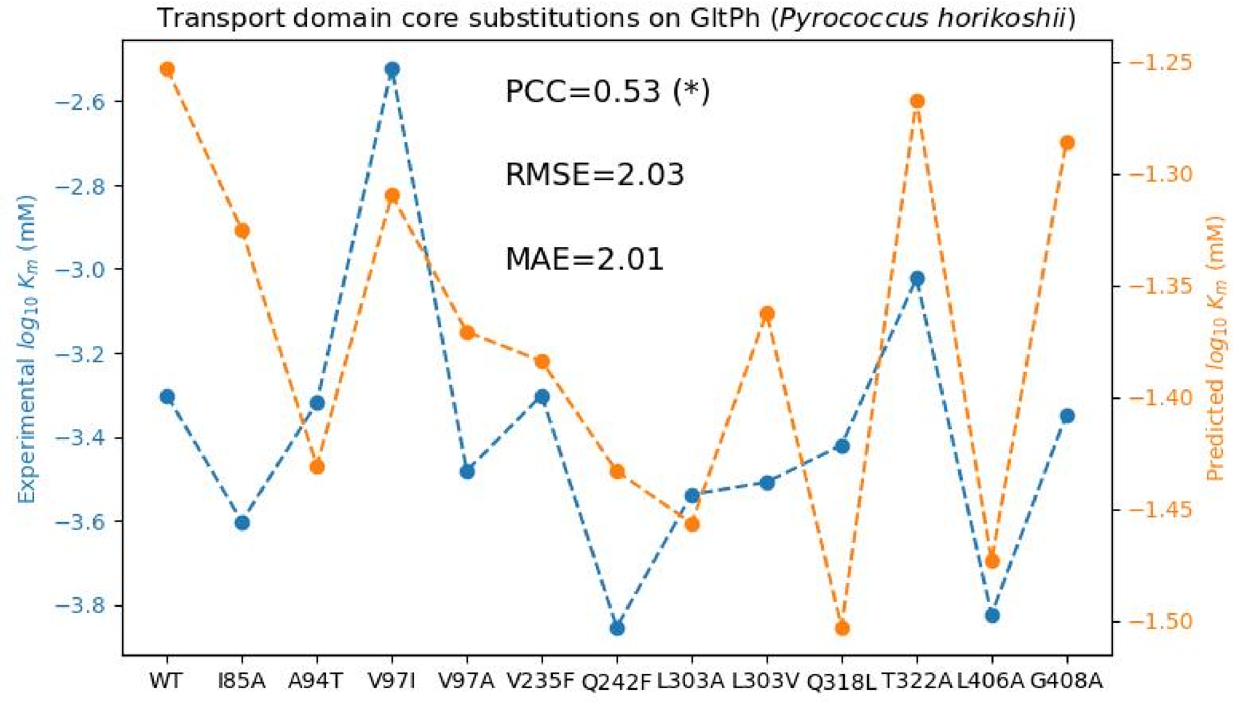


Figure S7. Experimental and predicted log10 $K_m$ of wild-type GltPh and GltPh with single-point mutations on the transport domain core.

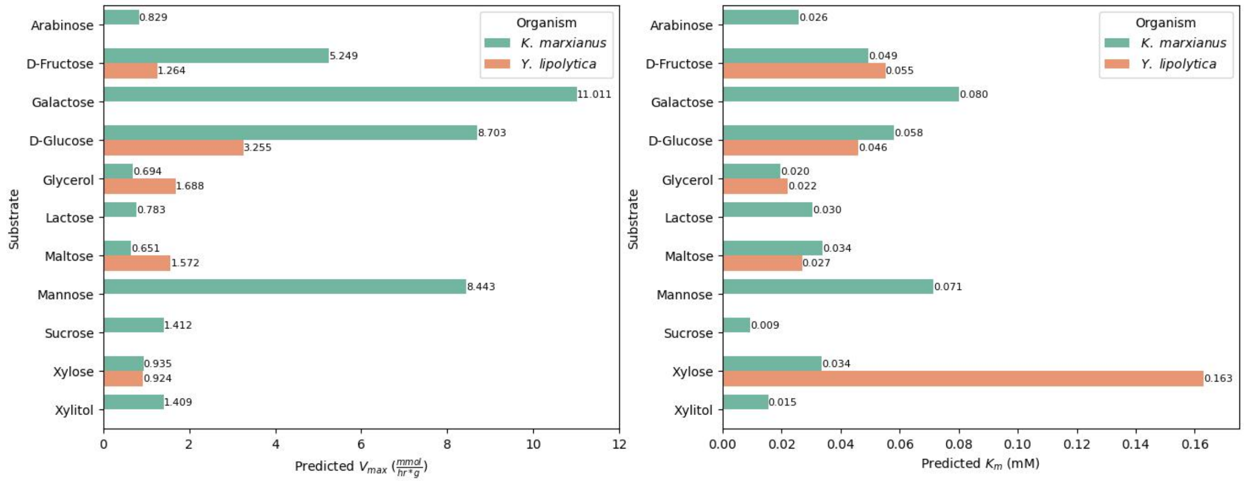


Figure S8. Predicted transporter $V_{max}$ (left) and $K_m$ (right) of *Y. lipolytica* and *K. marxianus* for different carbon sources.

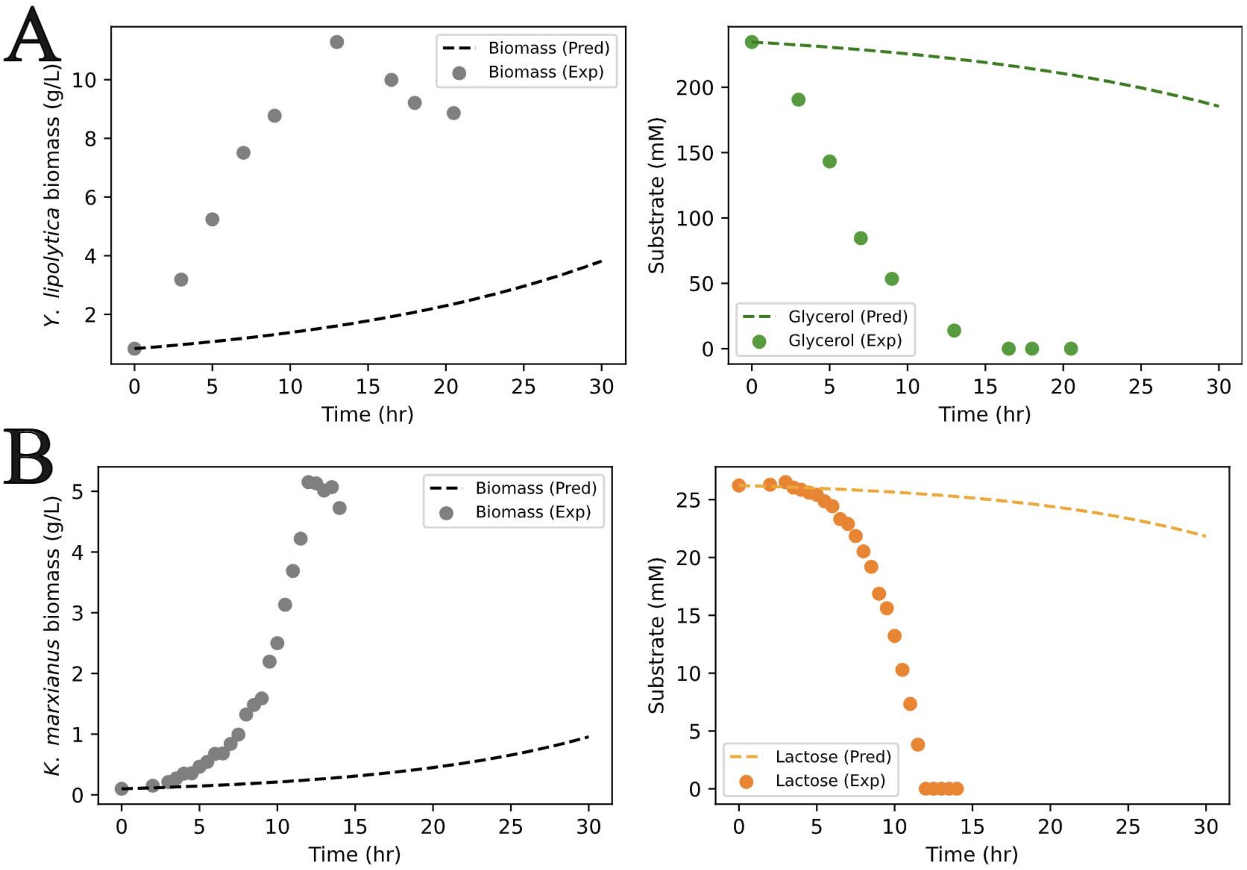


Figure S9. Dynamic TCFBA simulation of *Yarrowia lipolytica* using glycerol (A) and *Kluyveromyces marxianus* using lactose (B). Pred: predicted biomass or substrate concentrations, Exp: experimental biomass or substrate concentrations.

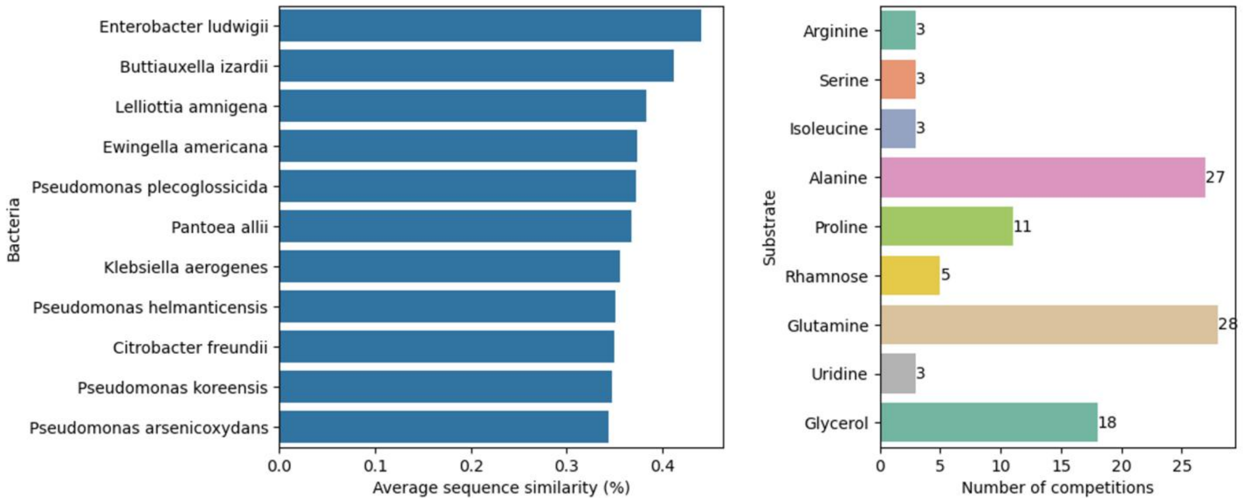


Figure S10. Supplementary information for the inter-species nutrient competition case study. (Left) Average sequence similarity scores of transporter proteins from 11 selected species, compared to the transporter sequences in the training datasets for $V_{max}$ and $K_m$. (Right) Numbers of inter-species competition relationships for 9 different substrates.

#### 2. Supplementary Tables

Table S1. Hyperparameter search space for XGBoost, LightGBM, ExtraTrees, and SVR

| Model | Search space |
|---|---|
| XGBoost | 1. booster: ['gbtree','dart']<br>2. n_estimators: 200~1000<br>3. max_depth: 3~10<br>4. learning_rate: 1e-3~1e-1<br>5. subsample: 0.5~1.0<br>6. colsample_bytree: 0.5~1.0<br>7. min_child_weight: 1~10<br>8. gamma: 0~5<br>9. reg_alpha: 1e-3~10.0<br>10. reg_lambda: 1e-3~10.0 |
| LightGBM | 1. n_estimators: 200~1000<br>2. boosting_type: ['gbdt', 'dart','rf']<br>3. learning_rate: 1e-3~1e-1<br>4. num_leaves: 16~256<br>5. max_depth: 3~10<br>6. min_child_samples: 5~50<br>7. subsample: 0.5~1.0<br>8. colsample_bytree: 0.5~1.0<br>9. reg_alpha: 1e-8~10.0<br>10. reg_lambda: 1e-8~10.0 |
| ExtraTrees | 1. n_estimators: 200~1000<br>2. max_depth: 5~12<br>3. min_samples_split: 2~20<br>4. min_samples_leaf: 1~10<br>5. min_weight_fraction_leaf: 0.0~0.05<br>6. max_features: 0.1~1.0 |
| SVR | 1. kernel: ["linear", "poly", "rbf", "sigmoid"]<br>2. C: 1e-3~1e3<br>3. epsilon: 1e-3~1<br>4. gamma: ["scale", "auto"]<br>5. degree: 2~5 |

Table S2. The best hyperparameter sets of top models

| Task | Model | Hyperparameters |
| --- | --- | --- |
| $V_{max}$ | XGBoost using Features (No FE) | 1. booster: 'gbtree'<br>2. n_estimators: 967<br>3. max_depth: 6<br>4. learning_rate: 0.0127636329718263<br>5. subsample: 0.6740195200487648<br>6. colsample_bytree: 0.600759202637599<br>7. min_child_weight: 6<br>8. gamma: 0.5846570301823268<br>9. reg_alpha: 0.0273900713673924<br>10. reg_lambda: 0.002030040853646 |
| $V_{max}$ | XGBoost using Features (No FE) + MFP | 1. booster: 'gbtree'<br>2. n_estimators: 967<br>3. max_depth: 6<br>4. learning_rate: 0.0127636329718263<br>5. subsample: 0.6740195200487648<br>6. colsample_bytree: 0.600759202637599<br>7. min_child_weight: 6<br>8. gamma: 0.5846570301823268<br>9. reg_alpha: 0.0273900713673924<br>10. reg_lambda: 0.002030040853646 |
| $K_m$ | XGBoost using Features (FE) | 1. booster: 'gbtree'<br>2. n_estimators: 770<br>3. max_depth: 3<br>4. learning_rate: 0.0399677393955049<br>5. subsample: 0.6987850532683515<br>6. colsample_bytree: 0.7585744899038346<br>7. min_child_weight: 5<br>8. gamma: 0.0248224558635815<br>9. reg_alpha: 0.0116291414945214<br>10. reg_lambda: 0.0018254260308112 |
| $K_m$ | LightGBM using Features (FE) | 1. n_estimators: 612<br>2. boosting_type: 'gbdt'<br>3. learning_rate: 0.0150658367105329<br>4. num_leaves: 131<br>5. max_depth: 10<br>6. min_child_samples: 17<br>7. subsample: 0.9943133516142176<br>8. colsample_bytree: 0.6124561670056058<br>9. reg_alpha: 3.682198196286383e-05<br>10. reg_lambda: 0.0069073427773744 |

Table S3. Transporter proteins for different carbon sources in *Yarrowia lipolytica* and *Kluyveromyces marxianus*

| Organism | Substrate | BIGG_ID | Transporter proteins (Uniprot_id) |
|---|---|---|---|
| *Yarrowia lipolytica* | D-glucose | glc__D | Q6CH11/Q6CG69/Q6C152/Q6CCJ1/Q6CCU6/Q6C152/Q6C4W0/Q6CG69/Q6CG30/Q6C8K6/Q6CAS4/Q6CI42/Q6C576/Q6CEA9/F2Z653/Q6CBQ5/Q6CDU0/Q6C2L7/Q6CAP1/Q6CFJ6/Q6C0E0 |
| *Yarrowia lipolytica* | glycerol | glyc | Q6C428/Q6C3E2/Q6C6W7/Q6C9U1/Q6C632/Q6C3A8 |
| *Yarrowia lipolytica* | xylose | xyl__D | Q6C4S4 |
| *Yarrowia lipolytica* | maltose | malt | Q6CH11/Q6CG69/Q6C4W0/Q6CG69/Q6CG30/Q6C8K6/Q6CAS4/Q6CI42/Q6C576/Q6CEA9/F2Z653/Q6CBQ5/Q6C2L7/Q6CAP1/Q6CFJ6/Q6C0E0 |
| *Yarrowia lipolytica* | D-fructose | fru | Q6C152 |
| *Kluyveromyces marxianus* | D-glucose | glc__D | W0T3I1/W0T533/W0T7U4/W0TCZ7/A0ABX6EX13/A0ABX6F2R3/A0ABX6EYX0/A0ABX6EVG2/A0ABX6EN84/A0ABX6EWH5/A0ABX6EN79/A0ABX6EPW4/A0ABX6EZ78/A0ABX6F014/A0ABX6F3U4 |
| *Kluyveromyces marxianus* | D-fructose | fru | W0TC80/W0TFY3/W0TDX2/W0TH43/W0TGD9/W0TBK0/W0TD85 |
| *Kluyveromyces marxianus* | lactose | lcts | W0T7D8/W0T8B1/W0TAG2/A0ABX6EZF5/A0A1T4IZL0/A0ABX6EW58/A0A1T4IZQ6/A0ABX6EXF2 |
| *Kluyveromyces marxianus* | galactose | gal | W0TC80/W0TFY3/W0TDX2/W0TH43/W0TGD9/W0TBK0/W0TD85/A0ABX6EX13/A0ABX6F2R3/A0ABX6EY82 |
| *Kluyveromyces* | xylose | xyl__D | W0T3I1/W0T533/W0T7U4/W0TCZ7 |

| | | | |
|---|---|---|---|
| *marxianus* | | | |
| *Kluyveromyces marxianus* | xylitol | xylt | W0T3I1/W0T533/W0T7U4/W0TCZ7 |
| *Kluyveromyces marxianus* | arabinose | arab__L | W0T3I1/W0T533/W0T7U4/W0TCZ7 |
| *Kluyveromyces marxianus* | mannose | man | W0T3I1/W0T533/W0T7U4/W0TCZ7/A0ABX6EX13/A0ABX6F2R3/A0ABX6F014/A0ABX6F3U4/A0ABX6EY82 |
| *Kluyveromyces marxianus* | maltose | malt | W0T3I1/W0T533/W0T7U4/W0TCZ7 |
| *Kluyveromyces marxianus* | sucrose | sucr | W0T3I1/W0T533/W0T7U4/W0TCZ7/A0ABX6EYX0/A0ABX6EVG2 |
| *Kluyveromyces marxianus* | glycerol | glyc | W0TDC5/W0TGF1 |
